# Changes in the urinary proteome in a Patient-Derived Xenograft model

**DOI:** 10.1101/365999

**Authors:** Yongtao Liu, Youzhu Wang, Zhixiang Cao, Youhe Gao

**Author notes:** Correspondence Youhe Gao, Department of Biochemistry and Molecular Biology, Gene Engineering Drug and Biotechnology Beijing Key Laboratory, Beijing Normal University, Beijing, 100875, China.

## Abstract

In this report, the urinary proteome from a patient-derived xenograft (PDX) model was compared at the peptide level to study the origins of urinary proteins in tumor-bearing nude mice. Urine was collected from the PDX mice before and after tumor implantation. A total of 515 mouse proteins were identified, of which 8 were differential proteins. Seventy-eight unambiguous human peptides from 42 human proteins were identified in the tumor-bearing group. Compared with the differential urinary proteins from the tumor-bearing immuno-competent rats, the differential proteins in the urine from the PDX model had no host immune response proteins in the very early stage urine in the tumor-bearing immuno-competent rat model.

## Introduction

Biomarkers are measurable changes associated with a physiological or pathophysiological process. Blood remains stable and balanced due to the homeostatic mechanisms of the body. In contrast, the urine is where most blood wastes are disposed, and thus, urine exhibits larger change (Gao, 2013).

Many reports found candidate biomarkers in the urine in the very early stage of disease. Pulmonary fibrosis-related proteins were detected before the formation of pulmonary fibrosis in the urine of a rat model, and if therapy could begin this timepoint, prednisone, which is ineffective in late stages, could effectively stop fibrosis development (Wu, 2017). In liver fibrosis model, urinary proteins underwent change earlier than aminotransferase and other indicators in the blood, and many of the changed proteins were associated with liver fibrosis, cirrhosis and their formation mechanisms (Zhang, 2017). In a multiple sclerosis model, changes in the urine proteome could be seen even before pathological changes occurred. Among the 7 proteins that changed the most, 6 were reported to be associated with multiple sclerosis (Zhao, 2017). The urinary proteins changed in a chronic pancreatitis rat model, and some of those differential urinary proteins were reported to be pancreatitis-related (Zhang, 2018). Several urinary proteins changed in a chronic obstructive pulmonary disease (COPD) rat model, and some of those candidate markers are COPD-associated (Huang, 2018). Many differential urinary proteins were detected in urine from a bacterial meningitis model, many of which are reported in cerebrospinal fluid and blood as biomarkers of bacterial meningitis (Ni, 2017).

The Walker 256 tumor cell line was subcutaneously injected into rats, and the urinary proteome changed significantly even before a tumor mass was palpable in the subcutaneous tumor model. Some of these proteins are reported as tumor markers or are associated with vaccine-related tumors (Wu, 2017). Similarly, C6 glioma cells were injected into rat brains and changes in the urinary proteome were found before detection by magnetic resonance imaging (MRI). Many of these differential urinary proteins were previously reported to be associated with glioma (Ni, 2017).

Since differential urinary proteins are present in the very early stages of cancer cell implantation, it is difficult to imagine that such a substantial change in the urine resulted from a small number of tumor cells. We wondered whether it was possible that theses urinary proteome changes resulted from the host response. In this study, we attempted to answer this using a patient-derived xenograft model (PDX model). As typical immunodeficient animals, nude mice have no normal T-cell immunity. Human tumor cells can easily grow into tumors in these mice. If most of the changes came from the implanted human tumors, human proteins should be unambiguously identifiable in the urine. If most of the early changes were from the host defense, some of the immune-related differential proteins in the nude mice would be missing compared to the differential urinary proteins from immuno-competent animals.

Intestinal tumor cells from patients were implanted subcutaneously into nude mice to construct the PDX model. Urine samples from nude mice were collected after tumors had formed. Urinary proteins were digested in-gel and profiled by LC-MS/MS.

## Materials and Methods

### Tumor acquisition and PDX model establishment

Fresh tumor samples of at least 6 mm × 6 mm × 6 mm (>200mm^3^) were obtained in situ or from metastatic colorectal cancer tissue after surgery. The sampling sites showed highly malignant tissue activity. After collection, the samples were repeatedly rinsed with precooled sterile saline and immediately placed in a precooled specialized preservative solution. The patient’s tumor tissues were transported to the laboratory at 4 °C and placed into a plate with RPMI-1640 medium (Gibco, USA). Tumor tissues were cleaned, and connective tissue, blood vessels, adipose tissue, calcification, and necrosis, were removed from the surface. Tumor tissues were selected and cut into 3 to 5-mm^3^ tumor blocks. Tumor growth in NOD/SCID inoculated at 4 subcutaneous points was observed daily. When the tumors reached a set volume, the tumor-bearing mice were sacrificed and the tumors dissected. Next, the tumors were cleaned, cut into small pellets of dimensions 3 mm × 3 mm × 3 mm, and inoculated subcutaneously into BALB-/-c-nu mice. Each mouse was inoculated at 1 point. After 23 days, urine was collected when the tumor volume reached 300 mm^3^.

### Urine collection and sample preparation

Animals were placed in metabolic cages overnight (for 8 h) individually to collect urine samples. During urine collection, mice had free access to water but no food to avoid urine contamination. Urine were stored at −80 °C and then centrifuged at 4 °C and 12,000 ×g for 30 min to remove cell debris. The supernatants were precipitated with three volumes of precooled acetone at −20 °C for 2 h followed by centrifugation at 4 °C and 12,000 ×g for 30 min. The precipitate was then resuspended in lysis buffer (8 mol/L urea, 2 mol/L thiourea, 25 mmol/L DTT and 50 mmol/L Tris)^[7]^. Protein concentrations were measured using the Bradford assay.

### Tryptic digestion

Urine samples from eight mice each from the control and tumor-bearing groups were selected and analyzed by mass spectrometry with in-gel digestion. Eighty micrograms of total protein was added to the preformed gel (Invitrogen, 8–12%, Carlsbad, CA), and the protein was separated by electrophoresis at 200 V, 40 min and dyed with Coomassie blue for subsequent easier to fading. Each gel was cut from the bottom up into 5 parts based on concentration, and each section was cut into 1 to 1.5-mm^3^ pellets. The pellets were washed with 25 mmol/L ammonium bicarbonate / acetonitrile (1:1 V/V) until the pellets were discolored. DTT was used at 20 mmol/L and 37 °C for 1 h to denature the disulfide bonds in the protein structure, and 55 mmol/L IAA was add for 30min in dark to alkylate the disulfide binding sites. Next, 5 ng/L trypsin (Trypsin Gold, Promega, Fitchburg, WI, USA) was added to the dried gel pellets and incubated at 37 °C overnight. Peptides were collected with 50 % acetonitrile, lyophilized and stored at −80 °C.

### LC-MS/MS analysis

Reconstituted peptides were desalted by C18 zip tip (Millipore, Germany) and dissolved in 0.1% formic acid. The Thermo EASY-nLC 1200 chromatographic system was loaded onto the precolumn and analytical column. Data were collected by the Orbitrap Fusion Lumos mass spectrometry system (Thermo, USA). The liquid chromatography method was as follows: precolumn: 75 μm x 2 cm, nanoViper C18, 2 μm, 100 Å (Thermo Fisher Scientific, USA), analytical column: 50 μm x 15 cm, nano Viper C18, 2 μm, 100 Å injection volume: 2 mL. The flow rate was 250 nL/min. Phase A was 0.1% formic acid/water (Fisher Scientific, Spain); phase B was 80% acetonitrile (Fisher Chemical, USA)/0.1% formic acid/20% water. The ion source was nano ESI, and MS data were collected by Orbitrap with a resolution of 120,000, ion charge range of 2–7, and 32% HCD. Secondary MS data were collected by Orbitrap with a resolution of 30,000.

### Label-free proteome quantification

Proteomic data were searched using the SwissProt *Homo sapiens* and *Mus musculus* databases (updated Sep 2017). Proteome Discoverer 2.1 software was used for processing. The search parameters were set as follows: the MS deviation of the peptide precursor and product ions was 0.05 Da; the ureido methylation was immobilized to cysteine, variable modifications were made to protein N-terminal acetylation and methionine oxidation, and two missing trypsin cleavage sites were allowed. The peptide FDR was less than 1%, and at least 2 specific peptides were identified pre protein.

### Bioinformatics analysis

Urine samples information for the nude mice was obtained from online freeware, DAVID Bioinformatics Resources 6.8 (https://david.ncifcrf.gov) for protein molecular functions, cell components, and biological processes.

## Results and Discussion

### Nude mice weights and tumor situation

The weights of nude mice in the tumor-bearing group increased slightly within days. As the size of the tumors increased, the net weight of the mice declined, but total weight remains the same, at an overall level of approximately 19–23g. The tumors in the tumor-bearing mice had a volume of approximately 300 mm^3^ when inoculated into BALB/c-nu mice for 23 days (**Figure 1**).

**Figure 1.**
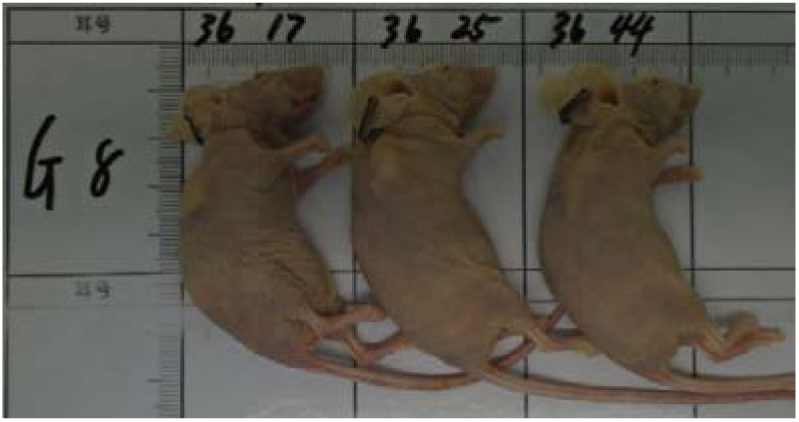
Nude mice in the tumor-bearing group. Tumor tissue was implanted subcutaneously in the right rear forelimb.

### SDS-PAGE analysis for urinary proteins

SDS-PAGE was used to compare urinary protein differences between the tumor-bearing mice and the control group (**Figure 2**). The differences were noted as followed: for the same sample amount, the tumor-bearing mice showed protein expressions of approximately 70 kDa, which was slightly increased, while the protein expression at 14 kDa decreased slightly. In general, no significant band appearance or disappearance was noted. From this SDS-PAGE result, 8 samples were selected from both the tumor-bearing and control groups, and their gels were extracted. Each gel was cut into 5 small samples, for a total 40 samples. After gel digestion, the proteins were broken down into peptides, and samples were analyzed by LC-MS/MS.

**Figure 2.**
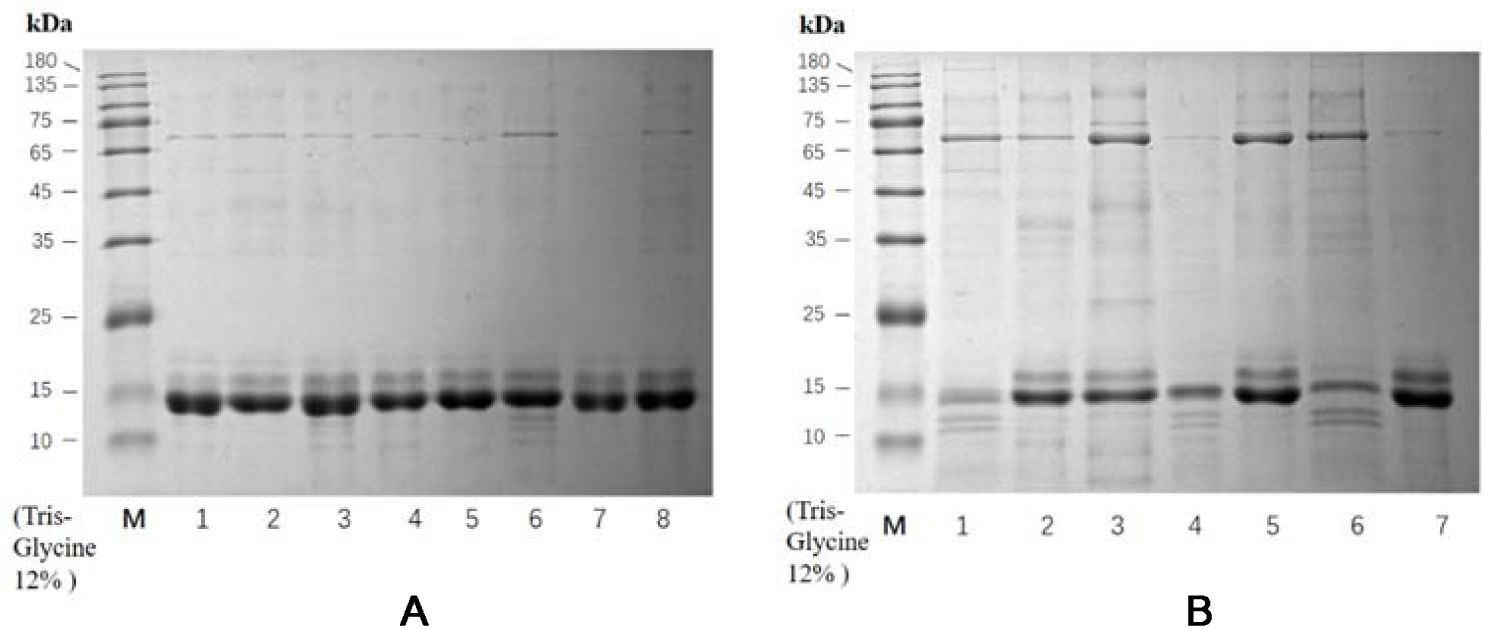
SDS-PAGE analysis of nude mouse urinary proteins. A: control group(n=8); B: tumor-bearing group (n=7).

### Urinary protein profiling from the PDX model

After processing by Protein Discoverer 2.1, the MS data were searched in the *Homo sapiens* and *Mus musculus* databases, to obtain protein names and peptide sequencing information. In total, 515 proteins were detected from the samples included both the control group and the tumor-bearing groups, and for each protein, at least 2 specific peptides were found. Two hundred four human proteins were detected, and after eliminating homologous proteins, 78 human-specific peptides and 42 proteins were obtained.

### Unambiguous human proteins

In the PDX model, when tumors were grown to 300 mm^3^, 42 unambiguous human proteins were found in the nude mice urine. Each human protein was searched in the human urinary protein database (https://www.urimarker.com/urine/) ^[9]^, which contains information on nearly 6,000 normal human urinary proteins. This database online and contains the most comprehensive information on human urinary protein biomarkers. By searching this database, protein abundances for each protein in normal human urine were determined.

Of the 42 human proteins, 21 were reported in normal human urine, of which 16 were high-abundance proteins (concentration greater than 1000 pg/mL), 3 were moderate-abundance proteins (concentration greater than 100 pg/mL), 2 were low-abundance proteins (concentration less than 100 pg/mL), and the other 21 did not appear in normal human urine. **Table 1** shows information on the numbers of total human proteins and specific peptides and their concentrations in normal human urine.

**Table 1.**
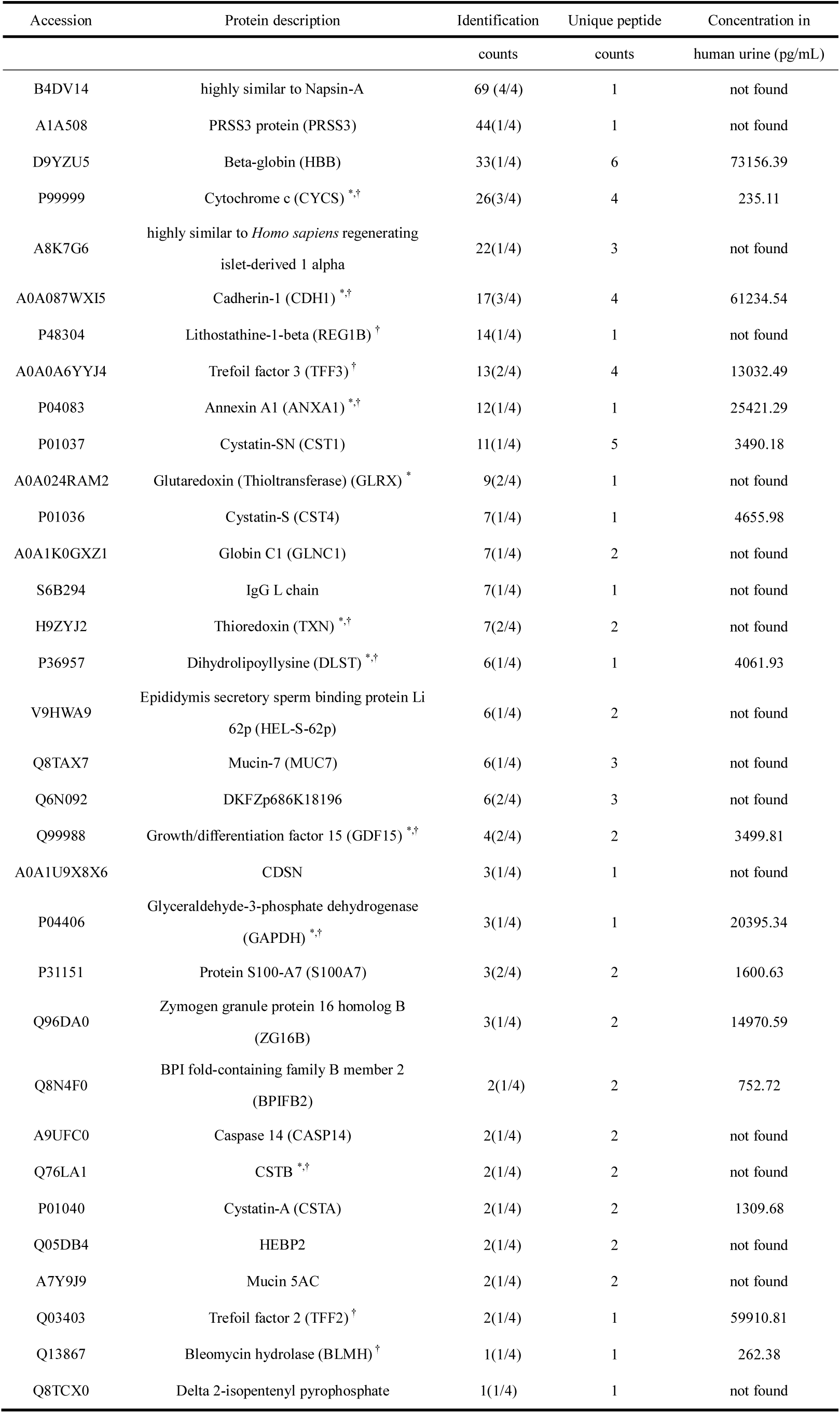
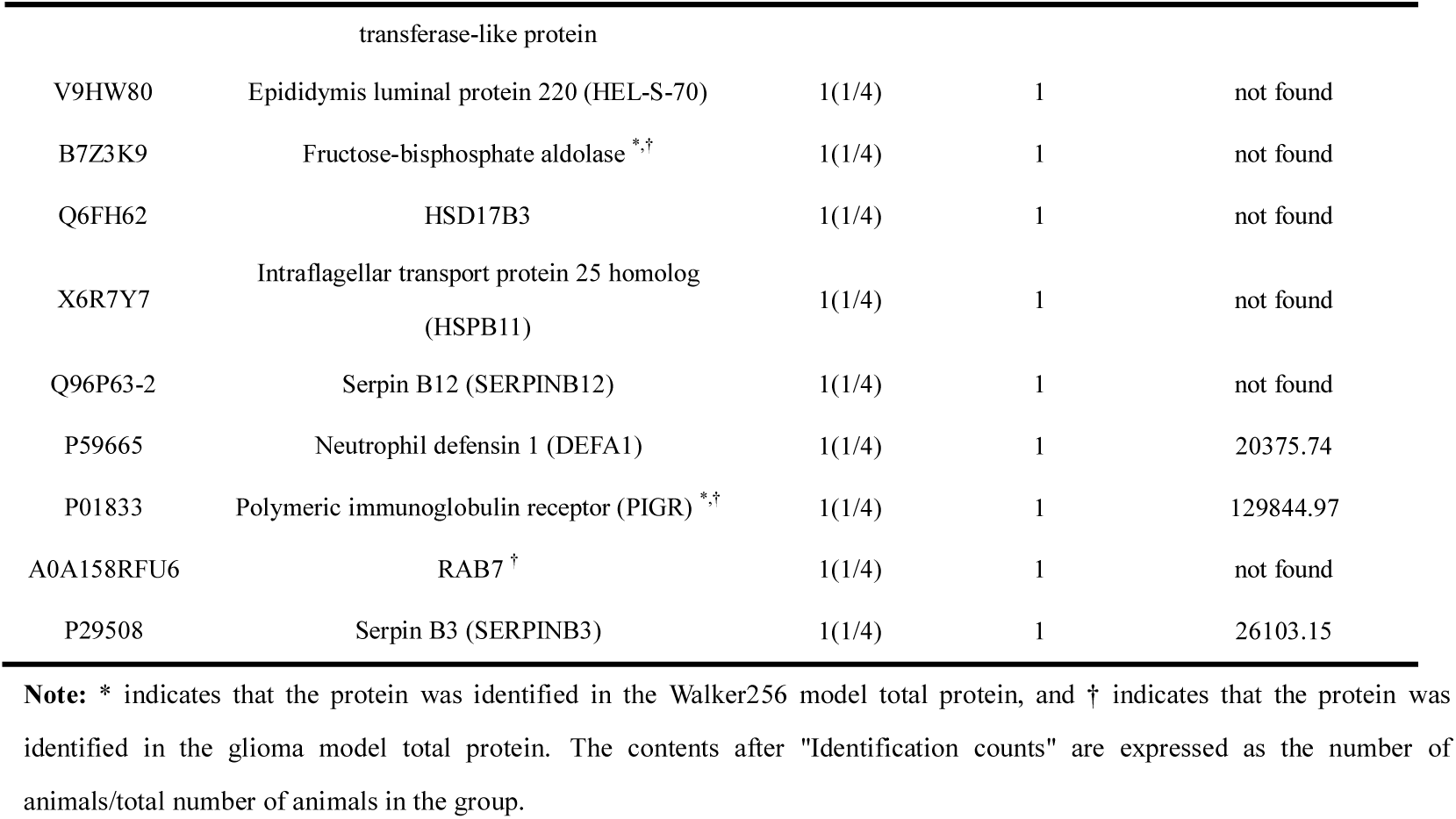
Human urinary protein information in tumor-bearing nude mice

This shows that in animals with normal immune systems, some differential urinary proteins were closely related to immune response. In immunodeficient animals, differential urinary proteins had little contact with the immune system.

The bioinformatics analysis results for the 42 human proteins are shown in **Figure 3.**

**Figure 3.**
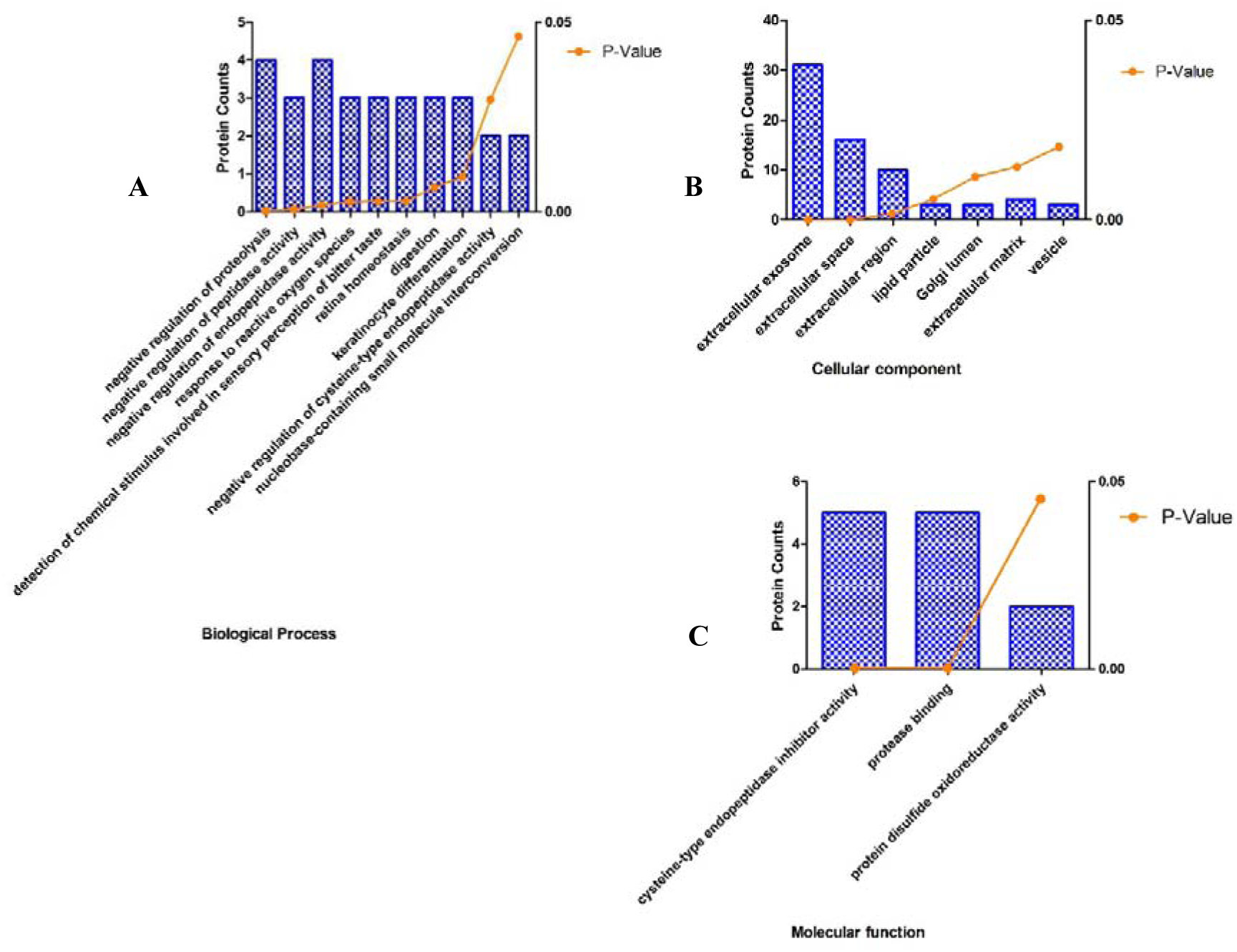
Cellular component (A); Molecular function **(B);** and Biological process **(C)** in the PDX human protein model.

### Differential urinary proteins in the host

The methods for screening differential proteins in mouse species are as follows. (1) Each protein contains at least two or more specific peptides. (2) After statistical analysis, the P value of each protein was less than 0.05 with a fold change is greater than 2. (3) The proteins were identified in each sample in the group (4/4), and 8 proteins met the above identification conditions. **Table 2** shows this information in detail.

**Table 2.**
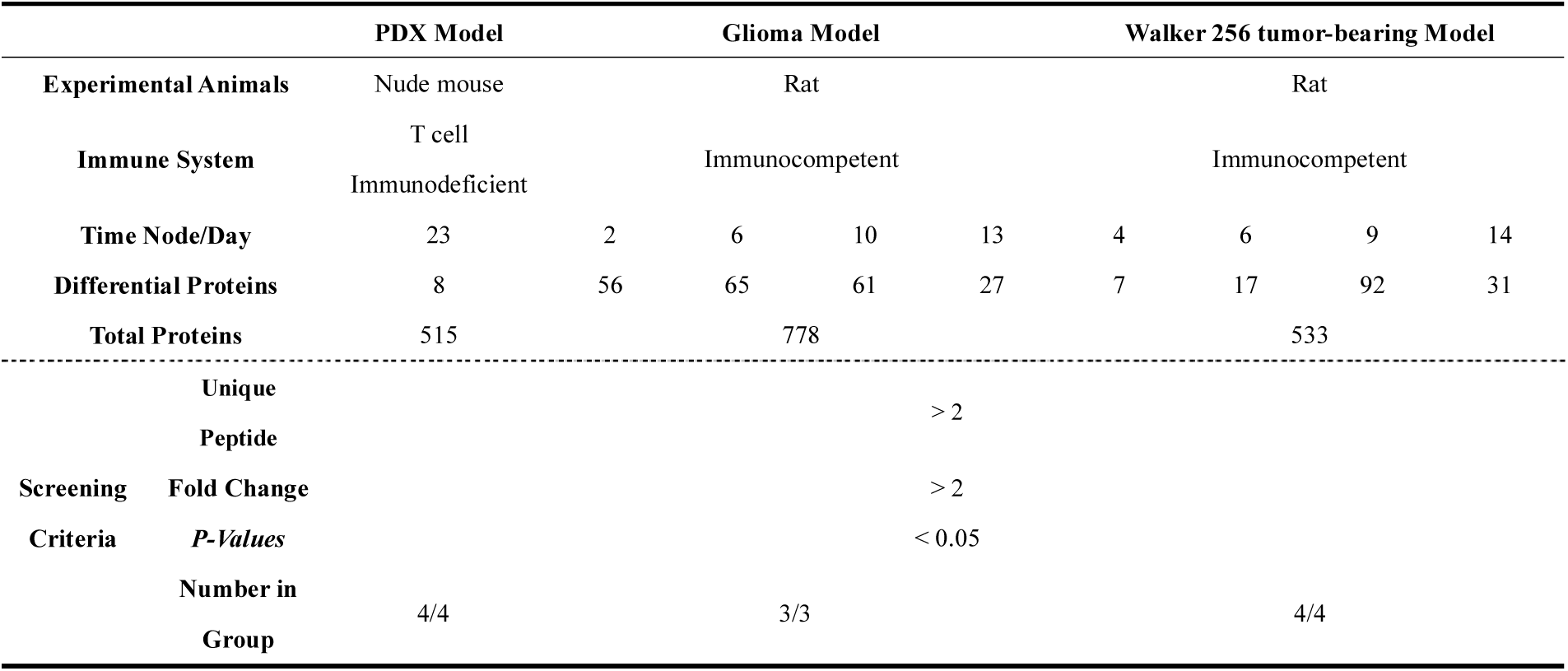
Differential urinary proteins in nude mice

The differential protein information obtained from these results was compared with Wu’s Walker256 rat model and Ni’s glioma rat model at different time nodes (**Table 3**).

**Table 3.**
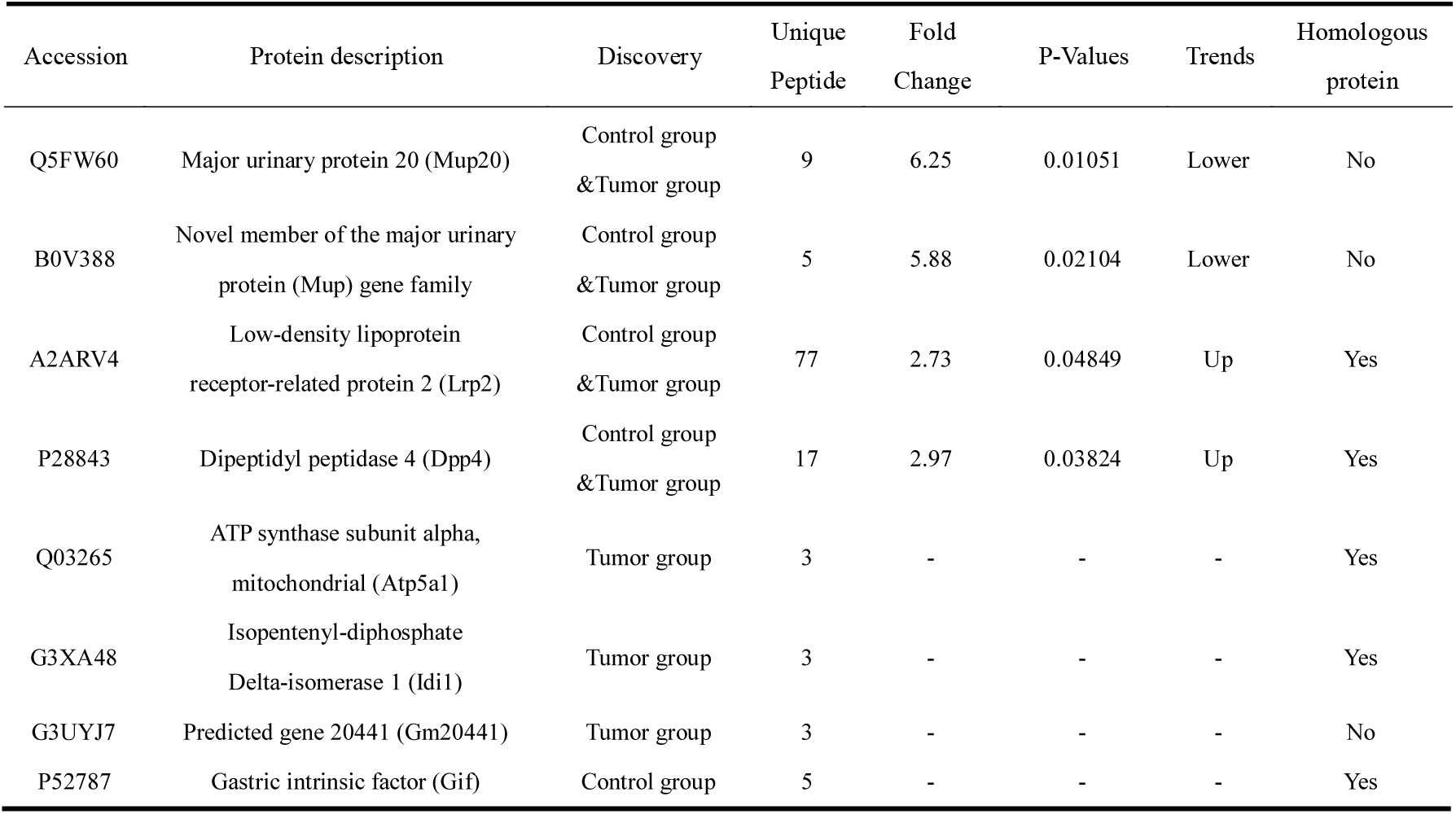
Comparisons of three tumor models and differential urinary protein information

### Urinary protein functional analysis

Biological process analyses were performed on 31 differential proteins on day 14 of the Walker 256 rat model and on 8 differential proteins of the PDX model. The 8 differential proteins of the PDX model were related to only the transport process (protein counts=3, p=0.083).

However, 32 biological processes were found in the Walker 256 rat model, many of which are related to immunity, such as complement activation, positive B-cell activation regulation, innate immune response, phagocytosis, the B-cell receptor signaling pathway, factor XII activation, and the immune system (**Table 4**). Compared with differential urinary proteins from tumor-bearing immuno-competent rats, the differential urinary proteins in the PDX model have none of the host immune response proteins found in the tumor-bearing immuno-competent rat model urine.

**Table 4.**
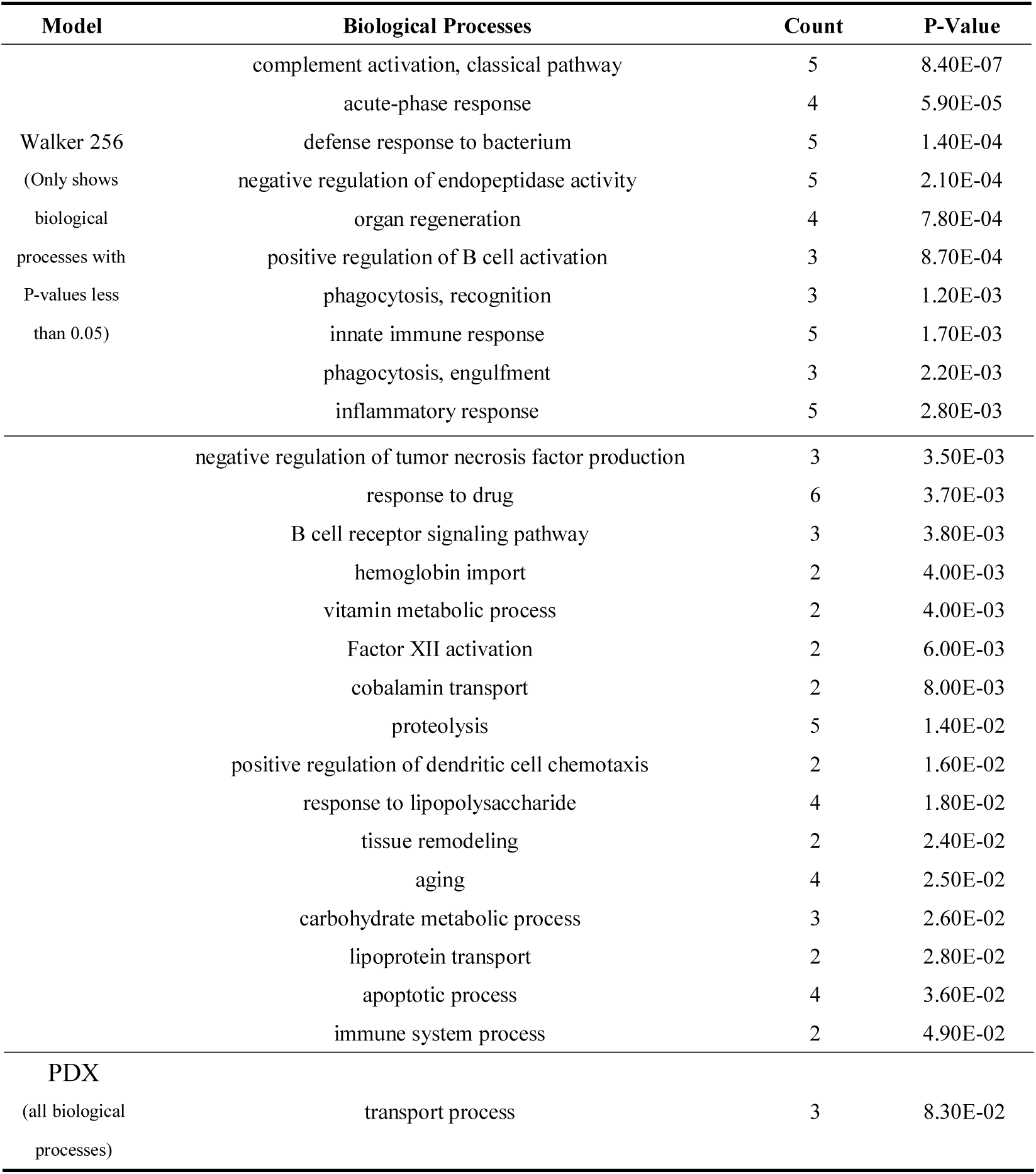
Biological process analysis on day 14 of the Walker 256 rat model and PDX models

Can the human proteins identified in the PDX be present in the urine of immuno-competent animals? Comparing 42 human proteins with the total urinary proteins identified by the Walker 256 and glioma rat models, 10 were identified in both models, 5 were identified in only the glioma rat model, and 1 was identified in only the Walker 256 model. The remaining 26 human proteins were not found in the total proteins of the other two models. This suggests that some human-origin tumor proteins identified in the PDX model may also be present in the urine of immuno-competent animals.

## Conclusion

Human-origin tumor proteins can be identified in PDX mouse urine. The differential urinary proteins in the PDX model had none of the host immune response proteins commonly found in the very early urinary stages in the tumor-bearing immuno-competent rat models.

